# Dual-Fluorescent Reporter for Live-Cell Imaging of the ER During DENV Infection

**DOI:** 10.1101/2022.09.08.507164

**Authors:** Lochlain Corliss, Madeline Holliday, Nicholas J. Lennemann

## Abstract

Infection by flaviviruses leads to dramatic remodeling of the endoplasmic reticulum (ER). Viral replication occurs within virus-induced vesicular invaginations in the ER membrane. A hallmark of flavivirus infection is expansion of the ER membrane which can be observed at specific time points post infection. However, this process has not been effectively visualized in living cells throughout the course of infection at the single cell resolution. In this study, we developed a plasmid-based reporter system to monitor flavivirus infection and simultaneous virus-induced manipulation of single cells throughout the course of infection in real-time. This system requires viral protease cleavage to release an ER-anchored fluorescent protein infection reporter that is fused to a nuclear localization signal (NLS). This proteolytic cleavage allows for the translocation of the infection reporter signal to the nucleus while an ER-specific fluorescent marker remains localized in the lumen. Thus, the construct allows for the visualization of virus-dependent changes to the ER throughout the course of infection. In this study, we show that our reporter was efficiently cleaved upon the expression of multiple flavivirus proteases, including dengue virus (DENV), Zika virus (ZIKV), and yellow fever virus (YFV). We also found that the DENV protease-dependent cleavage of our ER-anchored reporter exhibited more stringent cleavage sequence specificity than what has previously been shown with biochemical assays. Using this system for long term time-lapse imaging of living cells infected with DENV, we observed nuclear translocation of the reporter signal beginning approximately 8 hours post-infection, which continued to increase throughout the time course. Interestingly, we found that increased reporter signal translocation correlated with increased ER signal intensity, suggesting a positive association between DENV infection and ER expansion in a time-dependent manner. Overall, this report demonstrates that the FlavER platform provides a useful tool for monitoring flavivirus infection and simultaneously observing virus-dependent changes to the host cell ER, allowing for study of the temporal nature of virus-host interactions.

## Introduction

Flaviviruses belong to a large family of enveloped viruses with positive-sense single-stranded RNA genomes. The most prominent human viruses in this family are arthropod-borne and transmitted mainly through the bite of an infected mosquito (Valderrama et al., 2017; Weaver et al., 2018; Souza-Neto et al., 2019; Spadar et al., 2021). The prevalence of the arthropod vector of flaviviruses is the main contributor to the sporadic epidemic outbreaks in geographic locations with high mosquito populations. Members of this viral family cause a significant global health concern, with yellow fever virus (YFV) responsible for ∼ 200,000 cases per year and dengue virus (DENV) causing an overwhelming ∼ 390 million annual cases (Simo et al., 2019; Pierson and Diamond, 2020; Gaythorpe et al., 2021). Infection with flaviviruses can lead to disease resulting in severe clinical manifestations in multiple organ systems of the human body. Disease symptoms can include hemorrhagic fever or shock, caused by DENV and YFV, encephalitis, caused by West Nile virus (WNV), and birth defects such as microcephaly as a result of Zika virus (ZIKV) infection in pregnant women (Rossi et al., 2010; Duarte et al., 2017; Uno and Ross, 2018). Flavivirus epidemics have devastating effects on human health as well as significant social and economic burdens to the affected regions (Daep et al., 2014; Guo et al., 2017; Hameed et al., 2021). There are few safe and effective vaccines and no approved antiviral treatments for diseases caused by flaviviruses (Heinz and Stiasny, 2012; Ishikawa et al., 2014; Thomas and Yoon, 2019). Given the world-wide prevalence of flavivirus infections, it is crucial to continue the development of tools that can aid in obtaining a more in-depth understanding of the molecular processes of these pathogens that lead to manipulation of the host cell to advance the development of effective preventions and treatments.

The endoplasmic reticulum (ER) is the largest organelle of the cell and serves an array of important cellular functions, including production, trafficking, and degradation of proteins, metabolism of carbohydrates, and synthesis of lipids, making it an optimal site for flavivirus replication (Fagone and Jackowski, 2009; Braakman and Hebert, 2013; Ruggiano et al., 2014; Schwarz and Blower, 2016; Rajah et al., 2020). After viral entry of the host cell through endocytosis, the viral genome is directly translated as a single polyprotein that is embedded into the host cell ER through multiple transmembrane domains (Mazeaud et al., 2018; Ngo et al., 2019). The polyprotein is then co and post-translationally processed by viral and host proteases into ten individual subunits, composed of three structural proteins and seven nonstructural (NS) proteins (Amberg and Rice, 1999). The viral protease is made up of a complex between NS2B and NS3 (NS2B3). NS3 contains a serine protease domain that requires interaction with the ER-anchored NS2B protein via a cytoplasmic cofactor domain in order to be catalytically active (Falgout et al., 1991; Yusof et al., 2000; Xing et al., 2020). Thus, the active viral protease complex resides at the ER membrane in infected cells and is responsible for all cytoplasmic polyprotein cleavage events at a conserved motif, dibasic residues followed by Gly/Ser/Ala/Thr (Chambers et al., 1990, 1991; Shiryaev et al., 2007). Transmission electron microscopy has revealed remarkable manipulation of the ER during flavivirus replication, including the formation of replication vesicles that concentrate viral and host factors required for replication and packaging (Welsch et al., 2009; Gillespie et al., 2010; Paul, 2013; Junjhon et al., 2014; Nagy et al., 2016; Lennemann and Coyne, 2017; Arakawa and Morita, 2019; Evans et al., 2020). Several nonstructural proteins, such as the heavily membrane-associated NS4A and NS4B, have been shown to lead to drastic rearrangements of the ER membrane; however, expression of these proteins alone is not sufficient for the formation of replication vesicles (Kaufusi et al., 2014; Liang et al., 2016). Thus, viral manipulation of the ER is an important aspect during infection; however, the mechanisms and timing of this critical event are not well understood.

While the process of ER membrane remodeling has been studied utilizing the traditional method of imaging fixed cells, visualization of the events of virus-induced host cell manipulation throughout the course of infection has not been well-characterized. Imaging of fixed samples only allows for still snapshots of different cells at specific time points. However, the process of host cell manipulation during infection is very dynamic; therefore, still images cannot provide the full picture of the events that take place throughout the course of infection. To overcome these limitations, long-term time-lapse imaging of living cells infected with recombinant viruses expressing reporter fluorescent proteins has been used in previous reports (Schoggins et al., 2012; Shang et al., 2013; van der Schaar et al., 2016). However, genetic manipulation of the flavivirus genome to insert a reporter gene was shown to cause substantial attenuation of viral progeny. To bypass this obstacle, other studies have constructed plasmid-based reporter systems that rely on viral protease-dependent cleavage to release the fluorescent reporter signal to translocate to the nucleus and indicate infection (Jones et al., 2010; Medin et al., 2015; McFadden et al., 2018; Pahmeier et al., 2021). The limitation of these constructs is that they are able to detect only the presence of infection and cannot allow for the visualization of virus-induced changes to the host cellular structures. We previously developed a plasmid-based reporter to use in concert with live-cell imaging to detect infection by enteroviruses and track virus-induced manipulation of the host cell simultaneously (Lennemann et al., 2020). Therefore, we sought to expand on this construct by tailoring it to detect flavivirus infection and subsequent ER manipulation. In this study, we validate the efficacy of this reporter construct, termed FlavER, and uncover the temporal correlation between DENV infection and ER expansion.

## Results

### Flavivirus Reporter Plasmid Construction

To characterize flavivirus infection-dependent manipulation of host cell organelles in real-time, we modified our previously reported bipartite enterovirus reporter to include a flavivirus protease cleavage motif termed FlavER (Fig. 1) (Lennemann et al., 2020). It was hypothesized that this plasmid-based reporter system would allow for simultaneous monitoring of flavivirus infection and virus-induced changes to the ER. FlavER was constructed to express a cytoplasmic fluorescent protein fused to a simian virus 40 nuclear localization signal and the flavivirus NS2B3 protease cleavage sequence obtained through multiple sequence alignment of all cytoplasmic cleavage sequences processed by the viral protease within the DENV polyprotein (Fig. 1A). This fusion protein is anchored to the ER membrane via the transmembrane domain of the transferrin receptor (type II transmembrane domain) fused to an ER lumen localized fluorescent protein, which is fused to an ER retention signal (KDEL). Thus, we anticipated that the expression of the NS2B3 protease upon flavivirus infection results in cleavage of the consensus cleavage site, allowing for nuclear translocation of the ER-anchored infection reporter protein. Post cleavage, the fluorescent ER protein marker will remain in the ER lumen, allowing for visualization of changes to this organelle throughout the course of infection (Fig. 1B).

**Figure 1.**
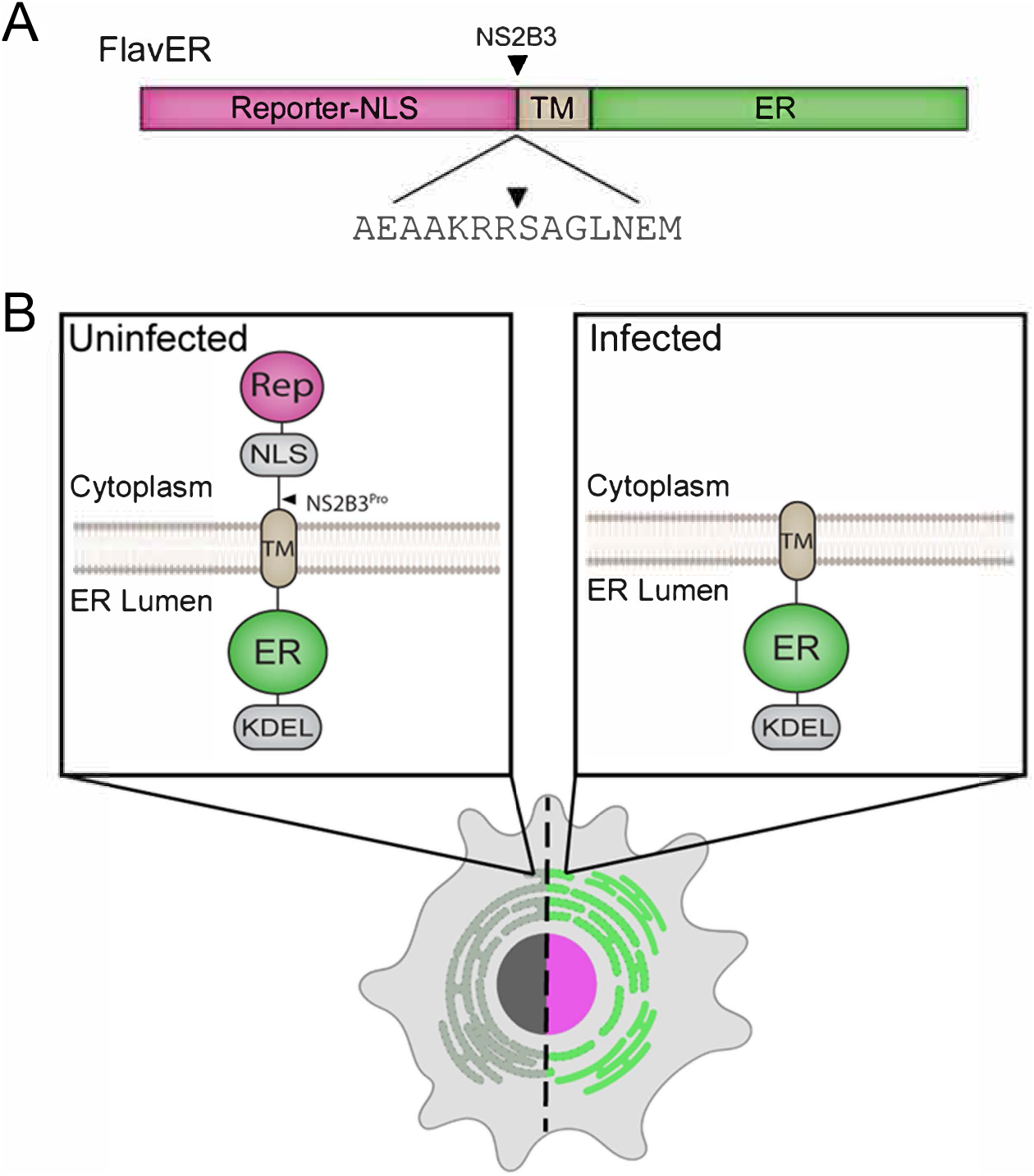
Design of an ER and flavivirus infection reporter. (A) Linear model of FlavER. The FlavER reporter consists of a fluorescent protein infection reporter fused to a nuclear localization sequence (NLS) followed by a consensus viral protease (NS2B3) cleavage sequence fused to a transmembrane domain (TM) followed by a fluorescent protein ER marker. Black triangle represents the site cleaved by the viral protease. (B) Proposed model of FlavER function. In uninfected cells (left), the reporter signal and ER marker colocalize at the ER. During flavivirus infection (right), viral NS2B3 cleavage releases the reporter signal, which then translocates to the nucleus.

### Validation of FlavER

To determine if FlavER could be cleaved by the dengue virus protease, U2OS cells were co-transfected with FlavER and V5 epitope-tagged active DENV NS2B3 (WT) or catalytically inactive DENV NS2B3 (S135A) and subjected to immunofluorescence (IF) staining or immunoblotting. IF staining of cells expressing FlavER and the active DENV NS2B3 showed localization of the protease-dependent reporter signal in the nucleus, while cells transfected with the catalytically inactive DENV protease (S135A) did not demonstrate this translocation (Fig 2A). Consistently, immunoblot analysis of cell lysates revealed that expression of the DENV NS2B-3 protease led to the production of a cleavage fragment representing the separation of GFP-NLS from the full-length FlavER construct. This cleavage fragment was absent in cells expressing the mutant DENV protease (Fig. 2B).

**Figure 2.**
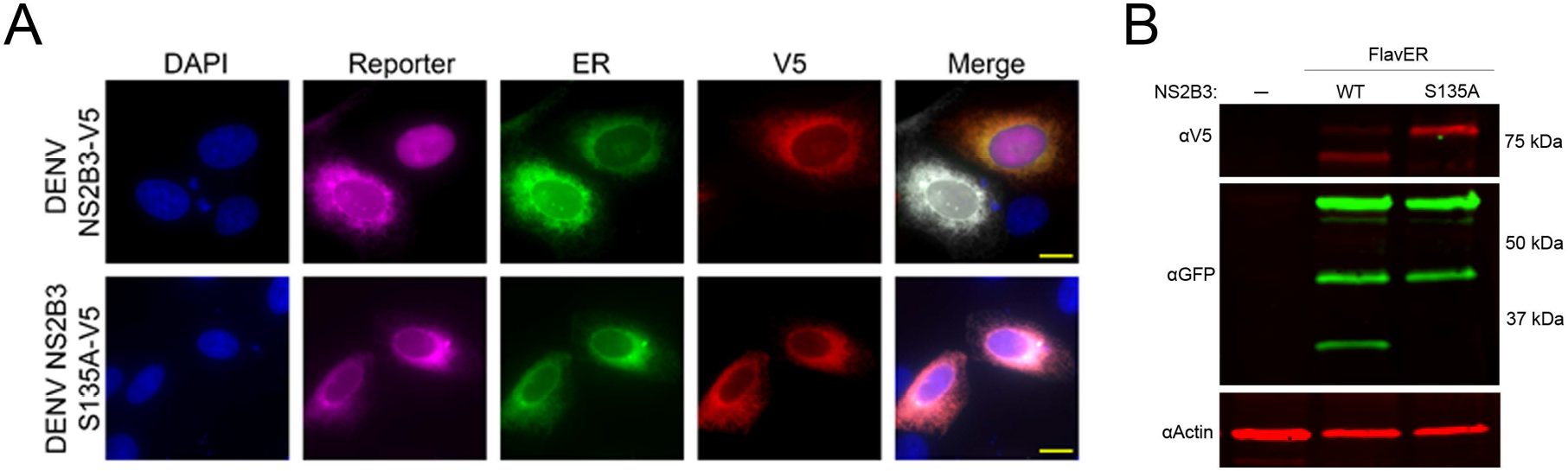
FlavER validation with DENV NS2B3. (A) Immunofluorescence (IF) microscopy of U2OS cells transfected with FlavER and an empty vector or V5-tagged DENV NS2B3. Infection reporter is shown in magenta, ER marker is shown in green, V5 staining is shown in red, and nuclei are stained with DAPI (blue). Scale bars represent 20 μm. (B) Immunoblot for V5, GFP, and actin of U2OS cells stably expressing FlavER transfected with an empty vector, active DENV protease, or inactive DENV protease (S135A).

We next wanted to test the efficacy of FlavER with DENV infection, so we first determined if FlavER expression resulted in viral attenuation. Titration of supernatants collected from WT and stable FlavER-expressing cells infected with DENV (0.1 FFU/cell) indicated no difference in viral production over 48 hours of infection (Fig. 3A). To assess the utility of FlavER during infection, we performed immunoblot analysis on lysates from DENV-infected cells expressing FlavER that were lysed at increasing hours post infection (Fig. 3B). We also analyzed lysates from FlavER-expressing cells that were infected with an increasing multiplicity of infection (MOI) of DENV (Fig. 3C). Both experiments revealed an increase of the cleavage fragment that was proportionate to the expression of DENV NS3 during the course of infection. Together, these results demonstrate the efficacy of FlavER in detecting protease activity upon expression of DENV NS2B-3 via transfection or infection.

**Figure 3.**
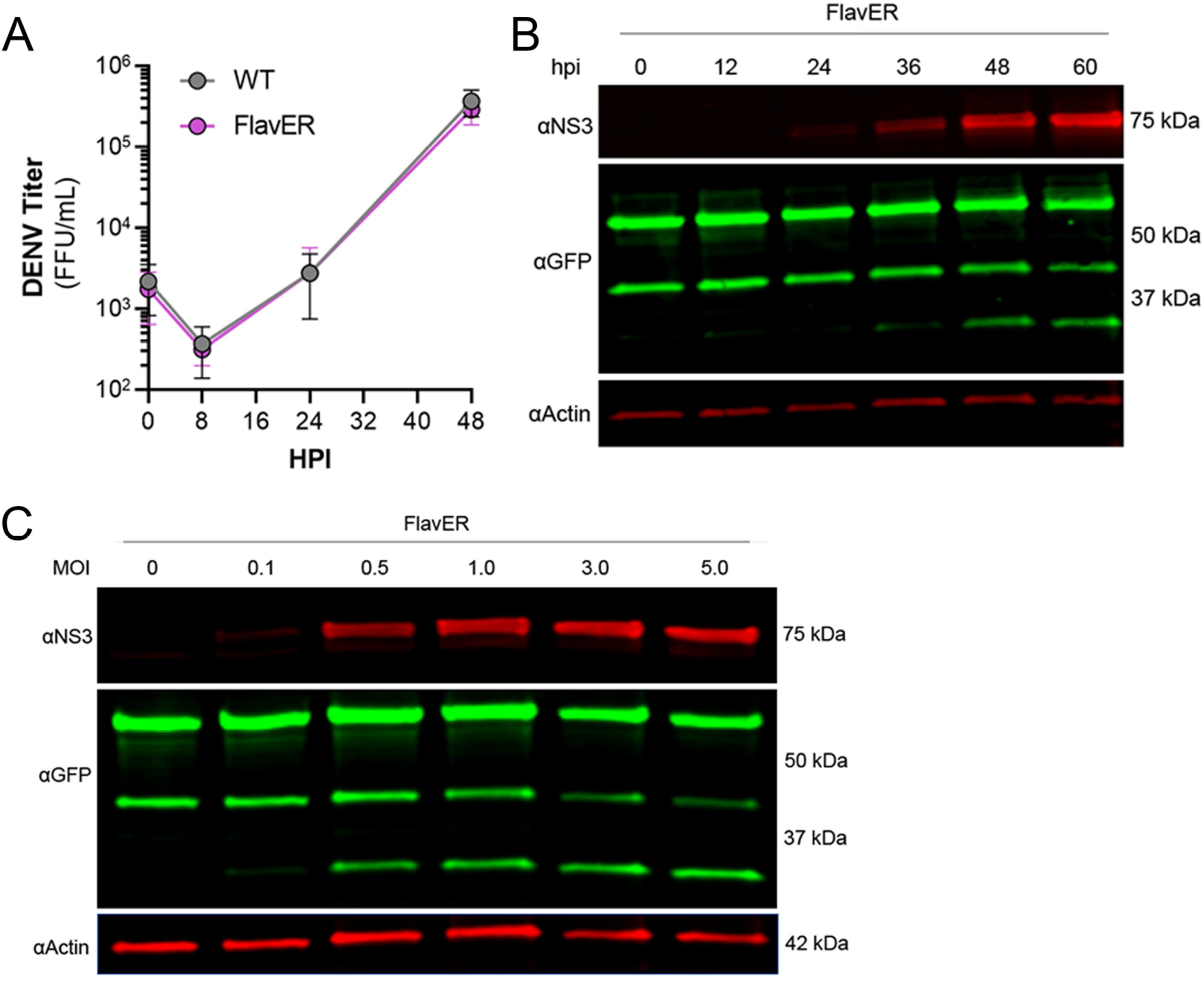
FlavER validation with DENV infection. (A) Growth kinetics of DENV in wildtype and FlavER-expressing Huh7 cells. Huh7 cells were infected with DENV at an MOI of 0.1, and FFU assays were performed on supernatants collected at the indicated time points, n=3. Data points represent the average titer (FFU/mL) for three independent experiments. (B) Immunoblot for NS3, GFP, and actin of U2OS cells stably expressing FlavER, infected with DENV at an MOI of 0.1, and lysed at the indicated hour post infection. (C) Immunoblot for NS3, GFP, and actin of U2OS cells stably expressing FlavER and infected with the indicated MOI of DENV.

### Multiple flavivirus proteases cleave FlavER

To determine if the consensus sequence in FlavER could be cleaved by multiple flavivirus proteases, we analyzed the lysates of FlavER-expressing cells transfected with the ZIKV, WNV, YFV, and mutant and active DENV proteases (Fig. 4A). Immunoblots showed FlavER cleavage by the DENV, ZIKV, and YFV proteases, but not by the WNV or mutant DENV proteases. We then quantified immunofluorescence images for the presence of the infection reporter in the nucleus of cells co-transfected with FlavER and DENV, DENV (S135A), ZIKV, WNV, or YFV NS2B3 (Fig. 4B). DENV, ZIKV, and YFV NS2B3 expression led to the efficient cleavage and release of the reporter signal with ∼82%, ∼79%, and 100% nuclear translocation, respectively. WNV NS2B3 was the least efficient in cleaving the reporter at ∼4% nuclear translocation, whereas no nuclear translocation of the reporter was observed with the catalytically inactive DENV NS2B3. Together, these data show the efficient cleavage of FlavER by DENV, ZIKV, and YFV proteases and the lack of cleavage performed by the WNV protease.

**Figure 4.**
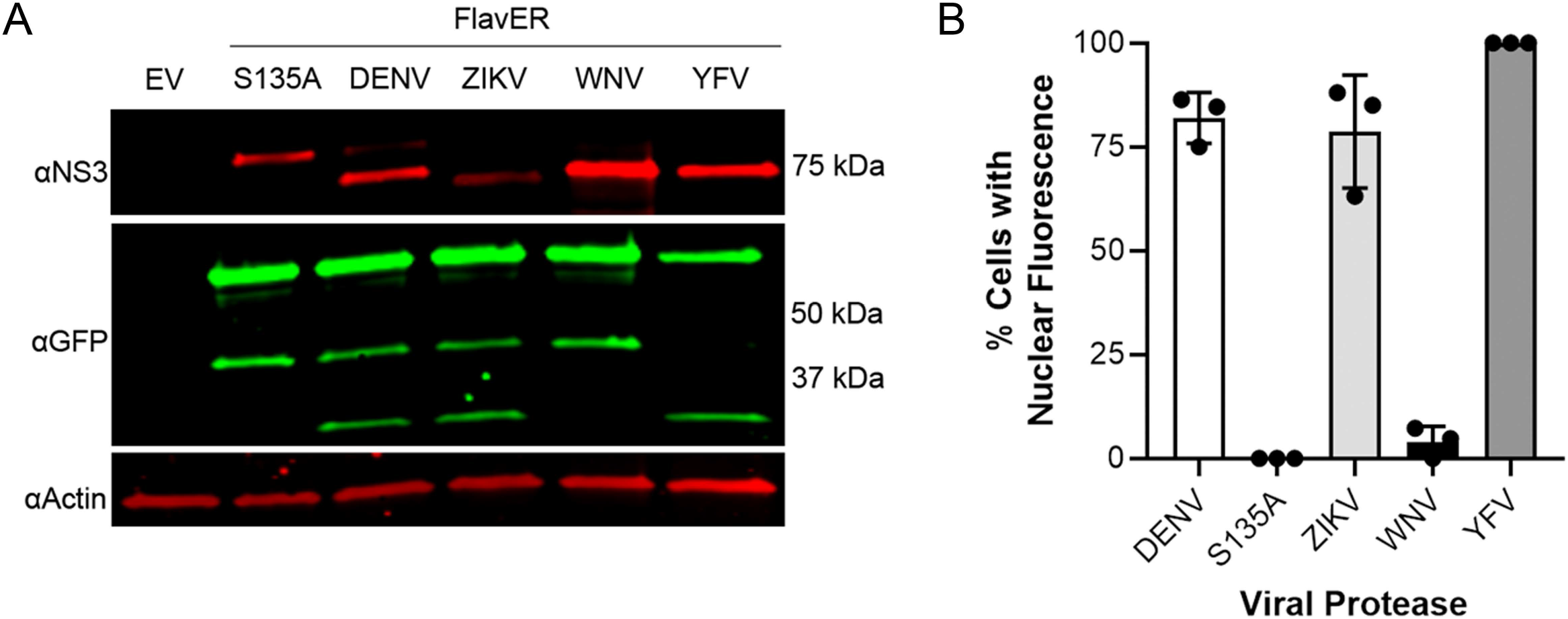
FlavER is cleaved by multiple flavivirus proteases. (A) Immunoblot for NS3, GFP, and actin for U2OS cells co-transfected with FlavER and the indicated flavivirus protease. (B) Quantification of IF images of cells expressing FlavER and the indicated V5-tagged viral protease. Data is presented as efficiency of cleavage determined by the percentage of co-expressing cells with nuclear-translocated fluorescent infection reporter, n=3. Data points represent results from three different fields of view, with the averages shown as bar graphs.

### DENV protease has more strict specificity for intracellular substrates

To better understand the applications of FlavER, we sought to determine how an intracellular reporter system differs from biochemical assays in terms of substrate specificity for the DENV protease. A previous study using biochemical assays with a soluble DENV protease and short peptide substrates showed that their protease was 90-100% effective at cleaving substrates with a phenylalanine and a proline as the residue positioned immediately after the dibasic residues in the cleavage motif (P1’), which are not naturally observed in the flavivirus polyprotein sequence (Shiryaev et al., 2007). We used site-directed mutagenesis to introduce single amino acid mutations to the P1’ site of our original FlavER construct, encoding a serine, to express a phenylalanine and a proline. U2OS cells expressing our WT and site-directed mutant reporters were infected with DENV at an MOI of 3 and lysed after 24 hours. Immunoblot analysis revealed that only the WT reporter with a serine in the P1’ position was capable of being cleaved by the DENV protease (Fig. 5A). Because all but one protease cleavages in the polyprotein occur proximal to the ER membrane, and the active protease localizes to the ER membrane, we sought to determine whether ER membrane-association of the substrate was a molecular determinant for cleavage. We constructed another mutant reporter from our original FlavER template where we removed the TM domain and the NLS of FlavER to make a soluble, protease-dependent cleavage reporter, termed FlavCyt due to its localization in the cytoplasm. U2OS cells were co-transfected with the active or mutant DENV protease and FlavER or FlavCyt. Immunoblot analysis of lysates showed that only the WT FlavER was able to be cleaved while FlavCyt was not, suggesting that reporter localization is a determinant for the ER membrane-associated DENV NS2B3 protease. (Fig. 5B). Together, these data highlight several determinants for studying the intracellular activity of the flavivirus protease.

**Figure 5.**
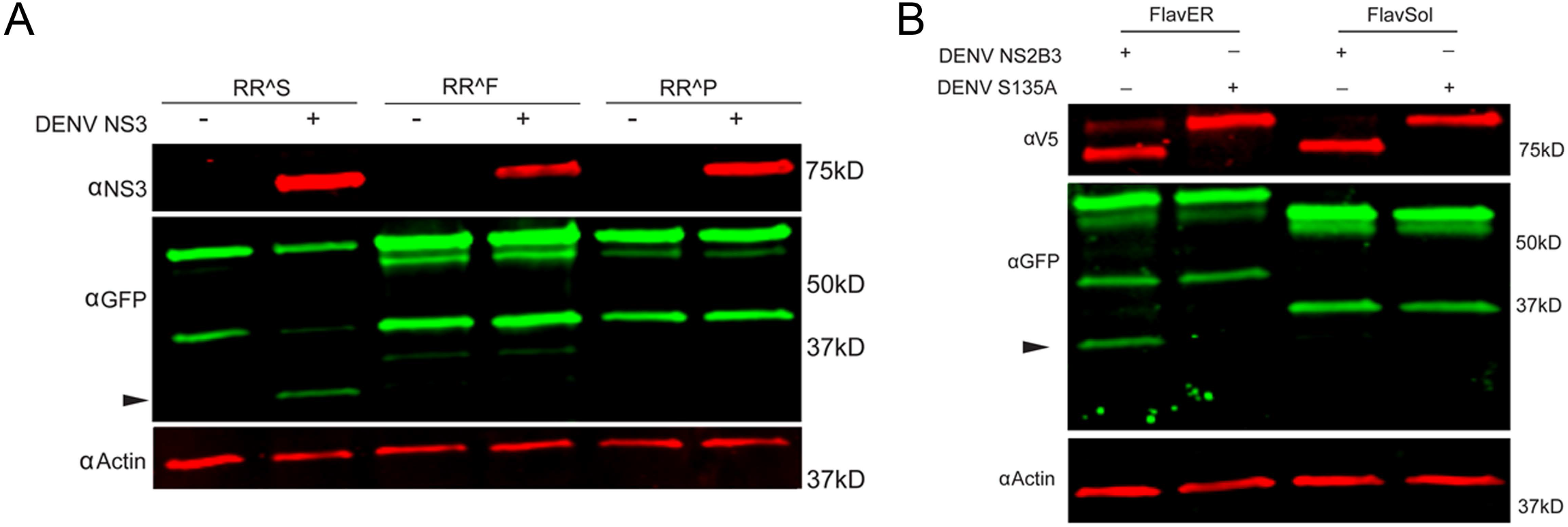
FlavER shows intracellular substrate specificity for the DENV protease. (A) Immunoblot for NS3, GFP, and actin for U2OS cells expressing the indicated FlavER construct and either left uninfected (mock) or infected with DENV at an MOI of 3. (B) Immunoblot of V5, GFP, and actin for U2OS cells expressing FlavER or FlavCyt and transfected with the indicated DENV protease.

### DENV infection leads to reporter translocation to the nucleus

To validate the efficacy of FlavER during infection, long-term time-lapse imaging was used to determine the efficiency of virus-induced nuclear translocation of the reporter signal. U2OS cells stably expressing FlavER were infected with DENV four hours prior to being placed in a humidified chamber set to 37°C for a 20-hour timespan with images taken in 20-minute intervals. DENV infection of cells expressing the reporter resulted in evident translocation of the fluorescent protein-NLS reporter to the nucleus, which was not observed in mock cells, while the ER-KDEL signal was shown to remain localized in the lumen throughout the course of a 24-hour infection (Supplemental Movie 1 and 2, Fig. 6A and B). Individual cells were quantified to determine the intensity of the reporter signal relative to the ER signal in the nucleus over time. Analysis of the image series showed that infected cells displayed a linear increase in nuclear-localized reporter signal throughout the course of infection (Fig. 6C). Uninfected (mock) cells exhibited no increase in nuclear reporter signal. Further analysis revealed that reporter translocation occurred in DENV-infected cells starting at ∼8 hours post infection, and 50% of the maximum signal was reached at ∼14 hours post infection on average (Fig. 5D). Taken together, these results suggest that the FlavER reporter can be used to detect infection via protease activity throughout the course of infection.

**Figure 6.**
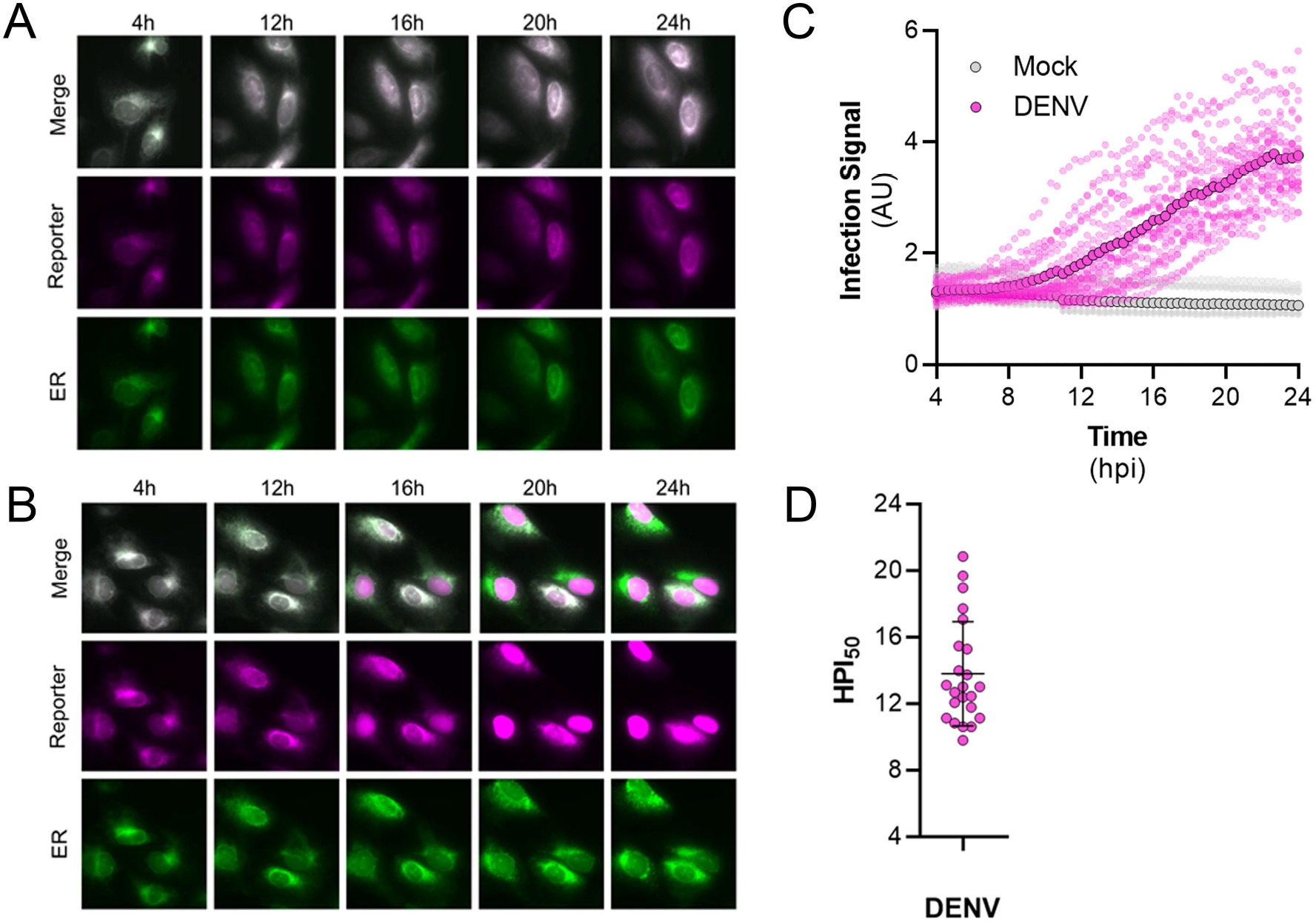
Time-lapse imaging of DENV infected cells expressing FlavER. (A,B) Representative time-points of an image series from live-cell imaging of the reporter signal (magenta), the ER marker (green), and merged panels from mock (A) or DENV infected (B) U2OS cells expressing FlavER. Scale bars represent 20 μm. (C) Quantification of infection intensity, defined as the ratio of reporter to ER fluorescence intensity signals within the nucleus from individual mock (gray, n=12) or DENV infected (magenta, n=23) U2OS cells expressing FlavER. Images were taken every 20 minutes, and infection intensity was calculated for every frame throughout a 20-hour time-lapse imaging experiment. Individual data points represent the average reporter intensity at the indicated timepoint. (D) Determination of the timepoint at which the 50% maximum reporter signal was observed in the nucleus (HPI50) as determined by nonlinear regression analysis for individual DENV-infected cells (n=23).

### DENV infection correlates with ER expansion in a time-dependent manner

To understand the kinetics of ER expansion as a result of infection by flaviviruses, we performed live-cell imaging of FlavER-expressing cells infected with DENV and quantified the resulting image series for changes in ER signal intensity over time. Cells infected with DENV showed a clear increase in the perinuclear ER signal intensity over the 20-hour imaging period, while uninfected cells showed no such increase (Fig. 7A). We also wanted to observe the relationship between infection and ER expansion, so we performed a correlation analysis which revealed that cells infected with DENV show a strong positive correlation between the infection reporter signal in the nucleus and perinuclear ER signal intensities (Fig. 7B). Together, these data show that as infection with DENV progresses, there is a quantifiable increase in perinuclear ER signal intensity.

**Figure 7.**
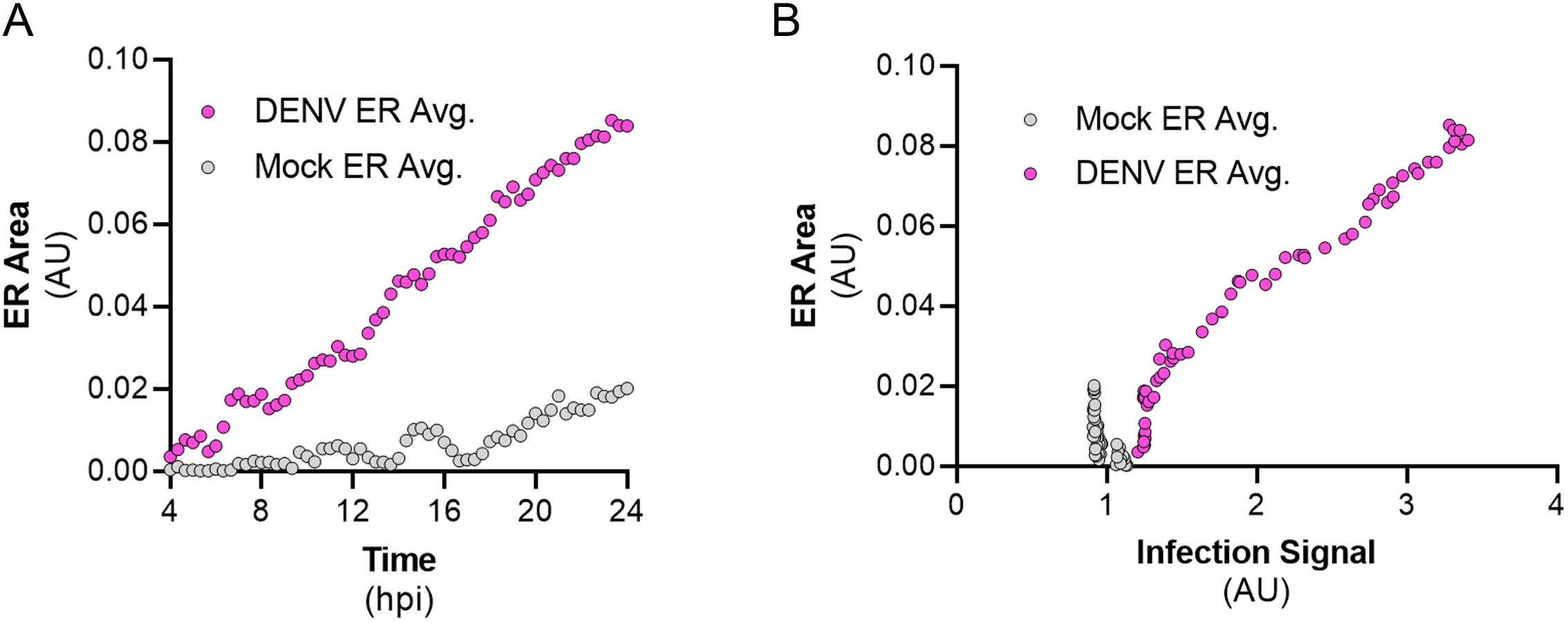
DENV infection correlates with ER expansion in a time-dependent manner. (A) Quantification of ER intensity for mock (gray, n=12) and DENV-infected cells (magenta, n=14) over the course of a 24-hour infection. (B) Correlation analysis between ER intensity and infection for mock (gray, n=12) and DENV-infected cells (magenta, n=14).

## Discussion

Long-term time-lapse imaging of living cells is a valuable method for studying virus-host interactions during the course of infection. In this study, this technique was used to monitor virus-induced changes to the ER of the host cell using a bipartite fluorescent reporter that requires flavivirus protease cleavage to release the NLS-tagged fluorescent protein. Release of this protein allows for its translocation to the nucleus of the host cell, while the ER-localized fluorescent marker is retained in the lumen. Additionally, we have shown that this system is a valuable tool for studying the determinants of flavivirus protease activity at the ER membrane, which is where it localizes during infection. Overall, this study shows that the design of the FlavER reporter provides a platform for understanding the temporal nature of virus-host interactions and protease activity during infection in real-time.

Previous groups have designed reporter viruses with a fluorescent tag inserted into the genomes of flaviviruses (Zou et al., 2011; Schoggins et al., 2012; Shang et al., 2013; Gadea et al., 2016; Suphatrakul et al., 2018; Tamura et al., 2018; Torres et al., 2022). These reporter viruses are useful in identifying infected cells; however, they do not accurately recapitulate the viral replication kinetics due to the attenuation that occurs when introducing a reporter gene into the genome. We have shown that our plasmid-based reporter system bypasses the issue of viral attenuation in cells expressing FlavER (Fig. 3A). Other studies have also employed plasmid-based reporter systems for live-cell imaging to monitor flavivirus infection in host cells. These groups designed their flavivirus infection reporters with the fluorescent reporter signal fused to viral non-structural protein 4B (NS4B) (Jones et al., 2010; Medin et al., 2015; McFadden et al., 2018). However, flavivirus NS4B is a multi-pass ER membrane-anchored protein that has been previously shown to induce autophagy, cause remodeling of the ER, and down-regulation of interferon-stimulating genes in cells expressing this protein alone (Munoz-Jordan et al., 2003; Kaufusi et al., 2014; Liang et al., 2016). These effects can make the results of experiments using these constructs difficult to interpret, which can be avoided using the FlavER system that has no impact on viral infection.

More recently, other groups have constructed plasmid-based reporters also dependent on viral protease activity; however, these reporters are limited to detecting infection and do not allow for the visualization of host structures unless additional markers are co-expressed in cells (Arias-Arias et al., 2020; Pahmeier et al., 2021). Previously, we designed a dual-fluorescent enterovirus reporter construct that allows for detection of infection and simultaneous visualization of host cell organelles (Evans et al., 2020). We then adapted this system to a single plasmid that can monitor flavivirus infection while observing the host cell ER in parallel. Therefore, FlavER improves upon previous constructs in that the design eliminates the overexpression of viral proteins known to manipulate the host cell and the need for transfection of multiple plasmids to simultaneously monitor infection and host cell manipulation. With the advantage of the ER-anchored fluorescent protein fused to the infection reporter, we were able to observe the changes that occurred in the ER throughout the course of DENV infection. Using live-cell imaging, we observed a quantifiable increase in ER signal intensity that positively correlated with the increase in infection (Fig. 7 A and B). This report uncovered the temporal nature of the relationship between DENV infection and virus-induced expansion of the ER. This correlation was unable to be shown previously with traditional methods of fixed cell microscopy, further demonstrating the usefulness of live-cell imaging to observe virus-induced manipulation of organelles at the single cell resolution.

Our reporter system not only allows us to monitor infection and ER manipulation in living cells but also provides a platform to investigate flavivirus protease specificity for intracellular substrates. First, we found that while the DENV, ZIKV, and YFV proteases are capable of cleaving the same consensus cleavage sequence, while the WNV protease had very poor cleavage efficiency, which is consistent with a previous report (Shiryaev et al., 2007) (Fig. 4B). These results suggest that different flavivirus proteases have distinct cleavage preferences for certain residues in intracellular substrates. Further, we determined that the DENV protease was unable to cleave versions of FlavER encoding Phe or Pro residues in the P1’ position (Fig. 5A). These results contradict what has been shown in previous biochemical studies, suggesting there are differences in protease activity in intracellular systems (Shiryaev et al., 2007). Additionally, our reporter system revealed that the presence of a transmembrane domain in the substrate may be another molecular determinant for flavivirus protease cleavage (Fig. 5B), which is supported by the ER membrane-proximal cleavage sites present in the viral polyprotein.

Overall, this study demonstrates the efficacy of our bipartite plasmid-based reporter to monitor viral infection and simultaneously observe virus-induced manipulation of the host cell in real-time, while also showing the importance of studying viral protease specificity in the context of the cell. Because this reporter construct allows for the indication of viral infection by distinct nuclear translocation of the infection signal to the nucleus, this system can be applied to high-throughput screens to identify antiviral therapeutics. By simply modifying the viral cleavage recognition sites, our reporter can be adapted for other positive-strand RNA viruses, including alphaviruses and coronaviruses. With the FlavER design, it is also possible to engineer the construct in a way that builds a multipartite reporter that allows for the visualization of virus-induced changes to multiple host cell structures. The reporter can be expanded for the observation of infection-induced changes to multiple host cell organelles simultaneously, such as the Golgi complex, mitochondria, lipid droplets, or the cytoskeleton by modifying the construct to contain three or even four fluorescent organelle marker proteins.

## Materials and Methods

### Cell Culture and Viruses

U2OS osteosarcoma cells (ATCC, HTB-96) and HEK-293T cells (ATCC, CRL-11268) were cultured in Dulbecco’s modified Eagle’s medium supplemented with 10% FBS and 100 IU penicillin/ 100 µg/mL streptomycin. Vero E6 cells (a gift from Dr. Kevin Harrod, University of Alabama at Birmingham) were cultured in modified Eagle’s medium supplemented with 10% fetal bovine serum and 100 IU penicillin/ 100 µg/mL streptomycin. C6/36 cells were cultures in Dulbecco’s modified Eagle’s medium supplemented with 10% FBS and 100 IU penicillin/ 100 µg/mL streptomycin. Human hepatoma cells (Huh7) were maintained in DMEM supplemented with 10% FBS, 9 g/L glucose, and 100U/mL P/S. All mammalian cells were maintained at 37 °C in a humidified environment with 5% CO_2_. C6/36 cells were maintained in a humidified environment at 28 °C.

Dengue virus serotype 2 strain 16681 (a gift from Carolyne Coyne, Duke University) was propagated in C6/36 cells (ATCC, CRL-1660) incubated at 33 °C for 6 days. Supernatants were combined with FBS to a final concentration of 20%, and cell debris was removed through centrifugation at 2300 xg for 15 minutes at 4 °C. Clarified supernatants were aliquoted and stored at -80 °C for future use.

### Focus Forming Assay

Viral stocks were titered by focus forming assay as previously described with minor modifications (Payne et al., 2006). Briefly, Vero cells were incubated with serial dilutions of dengue virus for 48 hours. Cells were fixed in 4% paraformaldehyde (PFA; Electron Microscopy Sciences) diluted in phosphate buffered saline (PBS; Corning, #21-040-CM) for 10 minutes at room temperature (RT), permeabilized in ice-cold methanol for 5 minutes at RT, and washed in PBS. Monolayers were incubated with α-flavivirus E-protein monoclonal antibody (clone 4G2, a gift from Margaret Kielian, Albert Einstein College of Medicine) for 1 hour at RT, washed with PBS, and incubated with secondary α-mouse antibody conjugated to Alexa Fluor 488 (Invitrogen, A11029) for 30 minutes at RT followed by a 5-minute incubation with 300 nM 4′,6-diamidino-2-phenylindole (DAPI; Invitrogen) diluted in PBS. Foci were counted using an Olympus IX83 inverted fluorescent microscope.

### Plasmid Construction

FlavER was cloned through the assembly of three DNA fragments into the NheI and XbaI restriction sites of pcDNA3.1(+) (Invitrogen). The first fragment was generated by amplifying the GFP from pcDNA3.1-GFP (a gift from Carolyn Coyne, Duke University) using Q5 high-fidelity DNA polymerase (New England Biolabs) with NheI-GFP_F (5’-GAAGCTAGCCACCATGGTGAGCAAGGGCGAGGAG-3’) and Xhol-GFP_R primers (5’-GAGACTCGAGCTTGTACAGCTCGTCCATGC-3’), followed by digestion with NheI-HF and XhoI (New England Biolabs). The second fragment was synthesized as a gBlock (Integrative DNA Technologies) consisting of 5’-XhoI – simian virus 40 nuclear localization sequence (- TCATCCGATGACGAGGCCACAGCTGATTCCCAGCACTCAACTCCGCCTAAAAAAAAAAGAA AAGTT) – KpnI – a flavivirus protease consensus cleavage sequence (GCAGAGGCTGCAAAAAGGAGGAGTGCTGGACTGAACGAGATG) – BamHI – transferrin receptor transmembrane domain (TATGGGACTATTGCTGTGATCGTCTTTTTCTTGATTGGATTTATGATTGGCTACTTGGGCTAT) – flexible linker (GGTGGATCTGGCGGAGGTTCCGGC) – EcoRI-3’, which was digested with XhoI and EcoRI-HF (New England Biolabs). The third fragment was generated by amplifying the mCherry-KDEL from pAc-mCherry-KDEL (a gift from Carolyn Coyne, Duke University) using Q5 high-fidelity DNA polymerase with EcoRI-mCh_F (5’- GTAGGGAATTCATGGTGAGCAAGGGCGAGGAGGATAAC-3’) and XbaI-mCh-KDEL_R (5’- GTAACGTTAGGGGGGGGGGATCTAGATCATAGCTCGTCTTTCTTGTACAGC-3’) to generate an EcoRI and XbaI flanked mCherry-KDEL. The three fragments were digested with the respective terminal restriction sites and ligated using T4 DNA ligase (New England Biolabs) for 3 hours at RT. The ligation was transformed into DH5α *E. coli* (Zymo Research).

pLJM1entr was generated to facilitate cloning of FlavER into pLJM1, a lentivirus transfer plasmid. The EGFP transgene in pLJM1-EGFP (a gift from was a gift from David Sabatini (Addgene plasmid # 19319, (Sancak et al., 2008) was replaced at the AgeI and EcoRI restriction sites with the following annealed oligos: Entr_F (5’- CCGGTTAATACGACTCACTATAGGGGATATCCCTCGACTGTGCCTTCTAG-3’) and Entr_R (5’- AATTCTAGAAGGCACAGTCGAGGGATATCCCCTATAGTGAGTCGTATTAA-3’) containing an EcoRV restriction sequence flanked by sequences homologous to the 5’ and 3’ untranslated regions of pcDNA3.1(+). pLJM1-FlavER was generated through the amplification of FlavER using Q5 high-fidelity DNA polymerase and pLJM1entry_F (5’- GAACCGTCAGATCCGCTAGCTAATACGACTCAC TATAGGG-3’) and pLJM1entry_R (5’-CATTTGTCTCGAGGTCGAGAATTCCACAGT CGAGGCTGATCAGC-3’) primers. This PCR product was assembled with EcoRV linearized pLJM1entr using HiFi DNA Assembly (New England Biolabs) according to the manufacturer’s instructions. This reaction was transformed into NEB Stable *E. coli*.

Expression plasmids for DENV NS2B-3-V5, DENV NS2B-3-S135A-V5, ZIKV NS2B-3-V5, and WNV NS2B-3-V5 have been previously described (Lennemann and Coyne, 2017). To construct YFV NS2B-3-V5, a plasmid expressing the yellow fever protease sequence was ordered from Twist Bioscience. This plasmid was then digested with NotI and KpnI restriction enzymes and ligated into the pcDNA3.1(+) vector expressing a V5 epitope tag.

### Immunofluorescence Microscopy

U2OS cells (40,000 cells/well) were reverse transfected with FlavER plasmid using Fugene HD (Promega) in an 8-well chamber slide (Celltreat). Twenty-four hours post transfection, cells were fixed in 4% PFA in PBS, permeabilized with 0.1% Triton X-100 (Fisher) diluted in PBS, washed with PBS, and incubated with mouse α -V5 epitope tag monoclonal antibody (Invitrogen, 46-0705) for 1 hour at RT. The monolayers were washed 3X with PBS for 5 minutes each and probed with goat α -mouse conjugated to Alexa Fluor-647 (Invitrogen, A21236) for 30 minutes at RT. The cells were washed 3X with PBS for 5 minutes each and incubated in PBS containing 300 nM DAPI for 5 minutes at RT. The slide was mounted using Vectashield Antifade Mounting Media (Vector Laboratories) and a 24×50mm Premium Superslip (FisherScientific). Three fields of view per well were captured using an Olympus IX83 inverted fluorescent microscope.

### Immunoblots

U2OS cells were transfected with the indicated plasmids using XtremeGene360 (Roche). When specified, cells were infected with DENV for the indicated timepoints. Cells were lysed in 1x RIPA + protease inhibitor + 0.1% SDS cocktail (Sigma). Lysates were sonicated prior to being separated by SDS PAGE using a 4-20% Tris-glycine polyacrylamide pre-cast gel (BioRad) and transferred to nitrocellulose membranes. Following 30 min of blocking in PBS + 10% non-fat milk, membranes were probed with the indicated primary antibodies: mouse anti-V5 (Invitrogen), rabbit anti-GFP (Proteintech), mouse anti-GFP (Proteintech), mouse anti-actin (Proteintech, rabbit anti-actin (Proteintech), and rabbit anti-DENV NS3 (GeneTex) followed by near-infrared dye-conjugated secondary antibodies (LiCor) diluted in PBST + 5% non-fat milk and imaged on an Odyssey CLx imaging system (LiCor).

### Lentivirus Production and Transductions

To produce lentiviral vectors expressing the FlavER transgene, HEK-293T cells (2 × 10^6^ cells/well) were reverse transfected with 1 µg pLJMI1-FlavER, 0.75 µg psPAX2 (a gift from Didier Trono (Addgene plasmid #12260), and 0.25 µg pCAGGS-G-Kan (a gift from Todd Green, University of Alabama at Birmingham) using Lipofectamine 3000 Transfection Reagent (Invitrogen) according to the manufacturer’s instructions. Lentivirus was harvested by passing the media through 0.2-micron filters (Thermo Scientific, 725-2520) 48 hours post transfection. U2OS cells (5 × 10^4^ cells) were reverse transduced with 50 µL FlavER lentiviral vector stock in growth media containing 10 µg/mL polybrene (Sigma Aldrich). Transduced cells were passaged for 48 hours and selected with 5 µg/mL puromycin for 5 days.

### Long-Term Time-Lapse Fluorescent Live-Cell Imaging

3 × 10^4^ U2OS cells stably expressing FlavER were transferred to an 8-well live-imaging slide (Ibidi). Cells were either left uninfected (mock) or infected with DENV at an MOI of 10. Infections were synchronized at 4 °C for 1h, followed by shifting cells to 37 °C for 3h prior to onset of imaging. One image per well was taken every 20 minutes over a 20-hour time period. Image series were cropped using ImageJ software (NIH, Bethesda, MA, USA), and multi-panel images were constructed using Photoshop CC 2021 (Adobe, San Jose, CA, USA).

### Image and Data Analysis

Models were made using Adobe Illustrator/Photoshop and Biorender. Percent nuclear translocation of the reporter signal in cells transfected with FlavER and a V5-tagged flavivirus protease was quantified using the counter plugin in ImageJ (NIH). The ImageJ counter plugin was also used to identify the number of the protease expressing cells with the reporter signal in the nucleus. Data was plotted using Prism 9 software (GraphPad, San Diego, CA, USA).

ImageJ was used to quantify relative infection signal from the live-cell time-lapse images. To account for non-specific fluorescence signal in the nucleus from the widefield microscope, the fluorescence signal intensity of the infection reporter was determined relative to the ER signal intensity within the same circular region of interest within the nucleus of individual cells from every image series. Data was plotted using Prism 9 software (GraphPad).

### Statistics

One-way ANOVA and nonlinear regression curve analyses were performed using Prism Software 9 software (GraphPad).

## Acknowledgements

We thank Carolyne Coyne (Duke University), Kevin Harrod (University of Alabama at Birmingham), and Todd Green (University of Alabama at Birmingham) for reagents. This project was supported by developmental funds from University of Alabama at Birmingham Department of Microbiology (NJL) and NIH grant K22AI143963-01 (NJL).

---

**Supplemental Movie 1**.

Live-cell imaging of individual DENV infected U2OS cells expressing FlavER. Movies were taken over a 20-hour timespan with images captured every 20 minutes.

**Supplemental Movie 2**.

Live-cell imaging of individual uninfected cells expressing FlavER. Movies were taken over a 20-hour timespan with images captured every 20 minutes.

## Notes

### Competing Interest Statement

The authors have declared no competing interest.

